# MicroRNAs and PFAS: A Pilot Study in Blood Collected from Firefighters

**DOI:** 10.1101/2024.04.05.588341

**Authors:** Xing Zhang, Mia Sands, Michael La Frano, Michael J. Spinella, Farzaneh Masoud, Christopher Fields, Zeynep Madak-Erdogan, Tor Jensen, Joseph Irudayaraj

**Affiliations:** Department of Bioengineering, University of Illinois, Urbana-Champaign, Urbana, IL 61801, USA; HPCBio, Roy J. Carver Biotechnology Center, University of Illinois, Urbana-Champaign, Urbana, IL 61801, USA; Department of Comparative Biosciences, University of Illinois, Urbana-Champaign, Urbana, IL 61801, USA; Illinois Fire Service Institute, University of Illinois, Urbana-Champaign, Urbana, IL 61801, USA; Carl Woese Institute for Genomic Biology, University of Illinois, Urbana-Champaign, Urbana, IL 61801, USA; Beckman Institute of Technology, University of Illinois, Urbana-Champaign, Urbana, IL 61801, USA; Cancer Center at Illinois, University of Illinois, Urbana-Champaign, Urbana, IL 61801, USA; Food Science and Human Nutrition, University of Illinois, Urbana-Champaign, Urbana, IL 61801, USA; Biomedical Research Facility, Carle Foundation Hospital, Urbana, IL 61801, USA

**Keywords:** PFAS, Firefighters, MiRNA, Exosome, NHANES

## Abstract

Per- and polyfluoroalkyl substances (PFAS) are chemicals with widespread industrial and consumer applications, and firefighters are known to be at risk of elevated PFAS exposure due to their occupational activities. This study aims to assess PFAS exposure and explore potential mechanistic insights through miRNA sequencing of plasma exosomes, in relation to PFAS levels in the general population. The study included 34 firefighter participants. PFAS levels in plasma were analyzed, and miRNA sequencing of plasma exosomes was conducted. The findings were compared with the general population data from the National Health and Nutrition Examination Survey (NHANES). While total PFAS levels did not significantly differ between firefighters and the general population in the cohort considered, variations in individual PFAS compounds were observed. MiRNA sequencing revealed substantial heterogeneity in miRNA expression patterns. Associations between serum PFAS levels and biochemical indicators suggested potential health implications, although further mechanistic insights need to be explored.

## 1. Introduction

Per- and polyfluoroalkyl substances (PFAS) are chemicals utilized across various industrial and consumer goods, primarily designed to provide resistance against heat, stains, water, and grease ^1^. Common exposure routes include Teflon, the coating on fast-food wrappers, non-stick pans, floor polish, carpets, upholstery fabrics, firefighting foams, treatments for clothing, and numerous other applications ^2^. Research indicates that these chemicals are present in the bodies of over 95% of Americans ^3^. Due to their exposure to smoke, the composition of firefighting foam, and the use of personal protective equipment (PPE) containing PFAS compounds ^4^, firefighters exhibit 3-5 times higher levels of circulating PFAS in their plasma ^5-9^. These substances persist in the environment for extended periods after release within and around active fire scenarios ^10^. This extended presence significantly heightens the potential for exposure in training facilities and firehouses, where firefighters spend substantial time ^11^. Research conducted through meta-analyses concentrating on the cancer vulnerability of firefighters has demonstrated an elevated likelihood for the development of several cancers, including multiple myeloma, non-Hodgkin lymphoma, prostate, kidney, lung, and testicular cancer ^12-14^. The increased susceptibility of firefighters to these cancers could stem from their direct exposure to intricate toxins during fire training exercises.

Exosomes are a class of vesicles of about 30-150 nm secreted by a variety of mammalian cells ^15^. Exosomes are currently regarded as a type of membrane vesicle capable of secreting specific components, which can affect the regulation of intercellular communication ^16,17^. Whether in normal physiological or pathological conditions, a variety of cells can produce and secrete exosomes which contain proteins, fats, mRNAs, and miRNAs ^18^. MicroRNAs are a class of endogenous non-coding RNAs (approximately 22 nucleotides) which regulate gene expression by base-pairing with target gene mRNA to degrade mRNA or hinder its translation ^19,20^. Under the regulation of multiple factors, especially the abnormally high expression of proto-oncogenes and the inhibition of tumor suppressor gene expression, tumor cells can acquire characteristics such as self-replication and immune escape ^21^. As an important post-transcriptional regulatory factor, microRNAs can participate in the regulation of tumor cell proliferation and migration ^22,23^. Exosomal miRNAs can be secreted into body fluids such as plasma, serum, gastric juice, tears, and urine, thereby avoiding degradation by ribonucleases ^24^. So far, studies have confirmed the value of miRNAs as biomarkers in the diagnosis and prediction of prognosis of patients with various tumors ^25-28^.

Current studies primarily examine associations of PFAS levels in firefighters’ blood with exposure levels and cancer risk, but only a few have performed mechanistic evaluation. For the first time, Jeong et al. examined miRNAs in the blood of firefighters and found that current firefighters exhibited differential microRNA expression compared with recruits, suggesting a potential mechanism for the development of cancer among firefighters ^29^. A longitudinal evaluation also supports this conclusion ^30^. Selecting participants from National Health and Nutrition Examination Survey (NHANES) and firefighters as research objects, it was found that there were significant negative associations among firefighters between PFAS level and interleukin-6 ^31^. Conversely, other studies showed that serum PFASs in firefighters are not associated with metabolic syndrome risk ^32^.

We hypothesized that PFAS exposure alters expression of miRNAs in exosomes and could be used as an indicator of PFAS toxicity and cancer risk assessment. To test our hypothesis, we collected blood samples from firefighters. Using GC-MS we quantified PFAS and metabolites in blood samples of two differentially exposed firefighter population. Finally, we conducted RNA-Seq analysis to assess the relationship between PFAS level in plasma and miRNA level in exosomes. We expect this study will provide a mechanistic basis for cancer risk evaluation or cancer biomarkers in populations exposed to PFAS.

## 2. Materials and Methods

### 2.1 Study Population of firefighters

Study participants were recruited (IRB #22226, OVCR, UIUC) from trainees at the Illinois Fire Service Institute (IFSI). New firefighters were recruited from basic fire-fighter academies that run in the Spring (April 2022) and Fall (September 2022). Experienced firefighters were recruited from weeklong onsite advanced firefighting classes, such as the Rapid Intervention Team (RIT) program, held throughout the year (November 2021-Septemebr 2022). A non-firefighter group was used as control. While discussing potential participation in the study, participants were provided with a detailed explanation of the purpose of the research and the potential risks of participation prior to obtaining informed written consent.

Participants were given the opportunity to ask questions. Based on exclusion criteria and previous history of recruiting subjects for similar studies ^8,33,34^, we screened approximately 50-60 firefighters to consent and enrolled 40 subjects. Inclusion criteria are firefighters 18-55 years old, a full-time employee (career and volunteer), a nonsmoker, and medically cleared by home fire department for participation in their occupational firefighting work. Exclusion criteria is firefighters over 55 or under 18 years of age, not a firefighter, currently a smoker/tobacco user, pregnant or lactating women.

### 2.2 Questionnaire

The questionnaire was conducted digitally through REDCap, a secure system designed for online data gathering and management. Physical measurements such as height, weight, and body fat index were collected from each participant prior to firefighting activity. Prior to participating in firefighting activities, participants completed the health history and physical activity questionnaire, which includes physical activities, typical PPE usage, cleaning, and exposure. The questionnaire is attached to the supplementary material S.4.

### 2.3 Plasma sample collection

Venous blood samples (10 ml) were collected in additive-free glass tubes by trained phlebotomists at the training site in IFSI and transported in ice for processing within 3 h of collection. The samples were subjected to centrifugation (3000 rpm) for 10 min after allowing for clotting at room temperature, and aliquoted into 1.2 mL cryo-vial tubes and stored at −80 °C until transport to University of Illinois Urbana-Champaign (UIUC) where they were processed.

### 2.4 Extraction of exosomes and miRNA

In this study, Invitrogen Total Exosome Isolation Kit (from plasma) (Catalog number: 4484450) was used to extract plasma exosomes from blood samples obtained from firefighters and the non-firefighter control group. About 200 μl of plasma from each sample was thawed and centrifuged at 2000×g and 10000×g for 20 minutes separately at room temperature to remove cell debris and large vesicles in the plasma. The supernatant was collected and diluted with PBS buffer and 10 μl of proteinase K was added and incubated at 37°C for 10 minutes to remove excess protein in plasma. 60 μl of exosome precipitation reagent was added and incubated at 4°C for 30 min. Next, the mix was centrifuged at 10000×g for 5 min and the exosome precipitate collected.

Total Exosome RNA & Protein Isolation Kit (Invitrogen) (Catalog number: 4478545) was used to extract miRNA in plasma exosomes for sequencing per manufacturer’s instructions ^35^.

### 2.5 Construction of strand specific RNAseq libraries

The DNA Sequencing Facility at the Roy J. Carver Biotechnology Center (University of Illinois at Urbana-Champaign) performed the library construction and sequencing studies using the Illumina NovaSeq 6000. To assess RNA integrity, total RNAs were processed through a Fragment Analyzer (Agilent, CA, USA). The TruSeq Stranded mRNA Sample Prep kit (Illumina, CA) was used to create RNAseq libraries. Enrichment of polyadenylated messenger RNAs (mRNAs) was performed using Dynabeads® Oligo (dT) beads from 500 ng of high-grade DNA-free total RNA. This step was followed by chemical fragmentation of the mRNAs and an annealing step with a random hexamer, and then conversion to double-stranded cDNAs, which are then blunt-ended, 3’-end A-tailed, and ligated to indexing adaptors. To avoid index switching, individual libraries were ligated to unique dual indexes. Using the Kapa HiFi polymerase (Roche, CA, USA), the adaptor-ligated cDNAs were amplified using 10 cycles of PCR. The resulting libraries are quantified using Qubit (ThermoFisher, MA, USA), and the mean library fragment length was calculated with a Fragment Analyzer. The libraries were further diluted to 10 nM and quantitated using qPCR on a Biorad CFX Connect Real-Time qPCR machine (Hercules, CA, USA) to ensure that barcoded libraries are correctly pooled and maximum clusters were in the flow cell. The pooled barcoded RNAseq libraries were placed onto a NovaSeq SP lane for clustering and sequencing. The sequencing of libraries was performed for 250 nt from either end of the fragments. The bcl2fastq v2.20 Conversion Software (Illumina, San Diego, CA, USA) was used to create and demultiplex the fastq read files.

### 2.6 Bioinformatics analysis of miRNA-seq data

Nf-core smrnaseq pipelines^36^ was used for sequencing alignment and the exploration of novel and known miRNA. (Nextflow version 22.10.6) ^37^. Initial quality control was done using FastQC ^38^ and 3’ adaptors were removed using Trim Galore ^39^. The resulting clean reads were then mapped to mature and hairpin miRNAs (miRBase v.22.1^40^), and GRCh38 human reference genome with Bowtie1 (v.1.3.1) ^41^. After alignment and trimming, sorted BAM files were used for further analysis with edgeR version 3.42.4 ^42^ and mirtop (http://mirtop.github.io). MiRNA quality was assessed and summarized in FastQC reports

### 2.7 Data sources and Study Population—National Health and Nutrition Examination Survey (NHANES)

NHANES (National Health and Nutrition Examination Survey - cdc.gov), an ongoing cross-sectional survey overseen by the Centers for Disease Control and Prevention was used. This survey employs a multistage cluster sampling method to ensure that its participants are a valid representation of the non-institutionalized population in the US. Individuals aged 12 and above are eligible, and blood samples were collected from them. Detailed information regarding the sampling procedure and data collection can be accessed on the NHANES website. For our study, we focused on investigating the relationship between per- and polyfluoroalkyl substances (PFAS) in serum and various biochemical indicators utilizing the existing survey data in this database. We selected individuals from the 2017–2018 NHANES cycles as our study population. We noted that in our chosen dataset of 9,254 subjects, only 8,704 completed both the interview and the Mobile Examination Center (MEC) examination within the specified cycle. Of these, one-third had their serum samples analyzed for a panel of 9 PFAS compounds. During our analysis, we addressed values falling below the lower detection limit by substituting them with a standardized value of 0.001. Subsequently, we aggregated the concentrations of these nine PFAS compounds to derive the total PFAS level for each participant.

### 2.8 Assessment of base model of general population and biochemical indicators from NHANES

We collected demographic information, lifestyle factors, and health conditions from the NHANES questionnaire to assess the base model. Demographic characteristics such as age, gender, race, education level, and family income-to-poverty ratio were included. Lifestyle characteristics, including Body Mass Index (BMI), smoking habits, alcohol consumption, and physical activity, were also considered. Furthermore, we examined various health conditions such as diabetes, hypertension, weak/failing kidneys, and cardiovascular disease. For our analysis, we utilized BMI as a proxy for measuring body weight and height. We categorized individuals into three smoking groups: non-smokers (with a smoking history of fewer than 100 cigarettes in a lifetime), current smokers (with a smoking history of more than 100 cigarettes and currently smoking), and former smokers (with a smoking history of more than 100 cigarettes but currently not smoking). Based on alcohol consumption, participants were grouped into non-drinkers (consuming no alcohol per day), moderate drinkers (consuming fewer than 2 drinks per day), and heavy drinkers (consuming 2 or more drinks per day). Physical activity levels were classified as low (exercising less than 2 times per week), moderate (exercising 2 to 5 times per week), and vigorous (exercising more than 5 times per week). Medical conditions were considered confirmed if respondents answered “yes” to questions regarding whether they had been told they had diabetes, hypertension, chest pain, or weak/failing kidneys.

To assess general health, we selected 26 biochemical indicators from the 2017-2018 NHANES laboratory results. These indicators are known to be representative of overall health, and existing literature suggests potential associations with per- and polyfluoroalkyl substances (PFAS). The selected biochemical indicators included albumin in urine, creatinine in urine, albumin in serum, creatinine in serum, ferritin and iron in serum, UIBC in serum, glycohemoglobin, folate in RBC, fasting glucose, total cholesterol, triglycerides in serum, ALT in serum, bicarbonate in serum, blood urea nitrogen in serum, chloride in serum, phosphorus in serum, potassium in serum, sodium in serum, total bilirubin in serum, total calcium in serum, total protein in serum, and uric acid in serum. We categorized each biochemical indicator into one of three groups, with the reference level being the normal range of bio-chemical indicators for the multivariate multinomial regression analysis.

### 2.9 Statistical Analysis

We incorporated sampling weights into all our analyses, in line with the NHANES sampling strategy. To assess the association of each biochemical indicator with categorical variables, we employed the Rao-Scott chi-square test. Similarly, to evaluate statistically significant differences among continuous variables, we utilized either the t-test for normally distributed data or the Mann-Whitney U test for non-normally distributed data (Supplementary Table S4). Our analysis proceeded in several steps. Initially, we applied multivariate linear regression to identify associations between log-transformed continuous weighted total serum PFAS levels and each biochemical indicator, adjusting for demographic factors, lifestyle variables, and health conditions. Subsequently, a multivariable multinomial logistic regression model was employed to determine the impact of serum PFAS levels on the likelihood of observing abnormal biochemical indicators. We computed odds ratios along with their corresponding 95% confidence intervals. In the multivariable analysis, we sequentially adjusted for covariates. Model 1 incorporated age, gender, race, education level, family income-to-poverty ratio, BMI, smoking habits, drinking habits, and physical activity. Model 2 further extended these adjustments to include specific health conditions: diabetes, hypertension, weak/failing kidneys, and cardiovascular disease. All statistical analyses were conducted using R version 4.3.0, and two-sided significance levels were calculated. We set the significance level at 0.05 for our analyses.

The information on chemicals and reagents (S.1), quantification of PFAS in blood (S.2), Quality assurance and quality control (QA/QC) (S.3), can be found in supplementary material.

## 3. Results

### 3.1 Participant Characteristics of firefighters

The characteristics of 34 firefighter (FF) participants divided between the new FF (N-FF, work experience less than 3 years) and experienced FF (E-FF, work experience more than 15 years), are presented in **Table 2**. Most enrollees were male (94.1%). The average age is 35.76 years old, and distribution of BMI index is relatively even. The average of N-FF experience is 1.36 years while the E-FF is 22.47 years. All FFs did not smoke, and most FFs drank alcohol (73.5%) and exercised more than 2 times a week (83.7%). A summary of demographic, lifestyle, and work exposure information is presented in Table S1 of the Supplementary Information. No sociodemographic and anthropometric data were significantly different between the two FF units.

**Table 1.** Firefighters’ Personal Protective Equipment statistics.

| Personal Protective Equipment | N-FF (n = 15) | E-FF (n=19) |
| --- | --- | --- |
| SCBA During Overhaul | 15 (100.0%) | 15 (78.9%) |
| % Time Wearing SCBA | 84.6% | 77.3% |
| Structural Knockdown - Coat/Pants | 15 (100.0%) | 19 (100.0%) |
| Structural Knockdown - Boots | 15 (100.0%) | 19 (100.0%) |
| Structural Knockdown - SCBA | 15 (100.0%) | 19 (100.0%) |
| Structural Knockdown - Helmet | 15 (100.0%) | 19 (100.0%) |
| Structural Knockdown - Hood | 15 (100.0%) | 19 (100.0%) |
| Structural Knockdown - Gloves | 15 (100.0%) | 19 (100.0%) |
| Structural Knockdown - Work Gloves | 12 (80.0%) | 6 (31.6%) |
| Structural Knockdown - Eye/Face Protection | 10 (66.7%) | 6 (31.6%) |
| Roof Ventilation - Coat/Pants | 15 (100.0%) | 19 (100.0%) |
| Roof Ventilation - Boots | 15 (100.0%) | 19 (100.0%) |
| Roof Ventilation - SCBA | 15 (100.0%) | 15 (78.9%) |
| Roof Ventilation - Helmet | 15 (100.0%) | 18 (94.7%) |
| Roof Ventilation - Hood | 15 (100.0%) | 18 (94.7%) |
| Roof Ventilation - Gloves | 15 (100.0%) | 19 (100.0%) |
| Roof Ventilation - Work Gloves | 10 (66.7%) | 6 (31.6%) |
| Roof Ventilation - Eye/Face Protection | 9 (60.0%) | 4 (21.1%) |
| Vehicle Fire Knockdown - Coat/Pants | 15 (100.0%) | 19 (100.0%) |
| Vehicle Fire Knockdown - Boots | 15 (100.0%) | 19 (100.0%) |
| Vehicle Fire Knockdown - SCBA | 15 (100.0%) | 18 (94.7%) |
| Vehicle Fire Knockdown - Helmet | 15 (100.0%) | 19 (100.0%) |
| Vehicle Fire Knockdown - Hood | 15 (100.0%) | 19 (100.0%) |
| Vehicle Fire Knockdown - Gloves | 15 (100.0%) | 19 (100.0%) |
| Vehicle Fire Knockdown - Work Gloves | 10 (66.7%) | 5 (26.3%) |
| Vehicle Fire Knockdown - Eye/Face Protection | 10 (66.7%) | 3 (15.8%) |
| Vehicle Fire Overhaul - Coat/Pants | 15 (100.0%) | 19 (100.0%) |
| Vehicle Fire Overhaul - Boots | 15 (100.0%) | 19 (100.0%) |
| Vehicle Fire Overhaul - SCBA | 11 (73.3%) | 14 (73.4%) |
| Vehicle Fire Overhaul - Helmet | 15 (100.0%) | 18 (94.7%) |
| Vehicle Fire Overhaul - Hood | 13 (86.7%) | 17 (89.5%) |
| Vehicle Fire Overhaul - Gloves | 14 (93.3%) | 19 (100.0%) |
| Vehicle Fire Overhaul - Work Gloves | 14 (93.3%) | 7 (36.8%) |
| Vehicle Fire Overhaul - Eye/Face Protection | 14 (93.3%) | 6 (31.6%) |
| Vegetation Fires - Coat/Pants | 14 (93.3%) | 19 (100.0%) |
| Vegetation Fires - Boots | 14 (93.3%) | 19 (100.0%) |
| Vegetation Fires - SCBA | 9 (60.0%) | 2 (10.5%) |
| Vegetation Fires - Helmet | 14 (93.3%) | 9 (47.4%) |
| Vegetation Fires - Hood | 13 (86.7%) | 9 (47.4%) |
| Vegetation Fires - Gloves | 13 (86.7%) | 11 (57.9%) |
| Vegetation Fires - Work Gloves | 13 (86.7%) | 13 (68.4%) |
| Vegetation Fires - Eye/Face Protection | 10 (66.7%) | 7 (36.8%) |

**Table 2.** Baseline characteristics of study population in NHANES 2017-2018 and firefighters.

| Characteristics | Overall<sup>1</sup> (n = 1048029) | PFAS<sup>2</sup> (n=1929) | Firefighters |
| --- | --- | --- | --- |
| Age | 34.4 ± 0.27 | 45.26 ± 0.47 | 35.76±1.76 |
| Gender |  |  |  |
| Male | 4273 (49.1%) | 952 (49.4%) | 32 (94.1%) |
| Female | 4431 (50.9%) | 977 (50.6%) | 2 (5.9%) |
| Race |  |  |  |
| Non-Hispanic White | 2931 (33.7%) | 667 (34.6%) |  |
| Non-Hispanic Black | 2010 (23.1%) | 430 (22.3%) |  |
| Non-Hispanic Asian | 1086 (12.5%) | 257 (13.3%) |  |
| Mexican American | 1298 (14.9%) | 297 (15.4%) |  |
| Other Hispanic | 773 (8.9%) | 176 (9.1%) |  |
| Others | 606 (7.0%) | 102 (5.3%) |  |
| Education level |  |  |  |
| Less than high school | 1052 (20.0%) | 329 (20.4%) |  |
| High school or equivalent | 1251 (23.8%) | 370 (23.0%) |  |
| College or above | 2950 (56.2%) | 912 (56.6%) |  |
| Family income-to-poverty ratio |  |  |  |
| ≤1 | 1797 (23.5%) | 358 (21.3%) |  |
| 1-3 | 3349 (43.9%) | 739 (44.0%) |  |
| >3 | 2488 (32.6%) | 581 (34.6%) |  |
| BMI |  |  |  |
| ≤25 | 3677 (45.9%) | 611 (32.1%) | 10 (29.4%) |
| 25-30 | 1920 (24.0%) | 556 (29.2%) | 13 (38.2%) |
| ≥30 | 2408 (30.1%) | 738 (38.7%) | 11 (32.4%) |
| Smoking |  |  |  |
| Current smoker | 972 (17.6%) | 311 (18.6%) |  |
| Former smoker | 1260 (22.8%) | 406 (24.3%) |  |
| Non-smoking | 3301 (59.7%) | 952 (57.0%) | 34 (100%) |
| Alcohol consumption |  |  |  |
| Non-Drinker | 572 (13.3%) | 162 (12.2%) | 9 (26.5%) |
| Moderate Drinker | 2988 (69.5%) | 936 (70.4%) | 5 (14.7%) |
| Heavy Drinker | 742 (17.2%) | 232 (17.4%) | 20 (58.8%) |
| Physical activity |  |  |  |
| Low | 2094 (37.9%) | 649 (38.2%) | 5 (14.7%) |
| Moderate | 1879 (34.0%) | 550 (32.4%) | 18 (52.9%) |
| Vigorous | 1554 (28.1%) | 499 (29.4%) | 11 (32.4%) |
| Self-reported diseases |  |  |  |
| Diabetes | 1028 (12.3%) | 304 (15.76%) | 2 (5.9%) |
| Hypertension | 2030 (34.9%) | 589 (32.8%) | 7 (20.6%) |
| Weak/failing kidney | 212 (4.0%) | 78 (4.8%) | 1 (2.9%) |
| Cardiovascular disease | 1086 (29.5%) | 589 (31.3%) | 1 (2.9%) |

<sup>1</sup> Overall population includes all interviewees who have also done the MEC exam. We only calculate the proportions of each category without the NA data.
<sup>2</sup> PFAS data only available from participants aged equal or greater than 12-year-old from a one-third sample population.

### 3.2 Comparison of the Distribution of DetectedPFAS

PFAS levels in plasma were measured by LC-MS. **Figure 1a** shows that there are no significant differences in total PFAS levels in plasma among the general population, N-FF and E-FF, and total PFAS value of most participants are between 10-30 ng/mL (One outlier is removed). Due to the high contribution, PFHxA, PFHxS and PFOS are analyzed separately. Results indicated that PFHxA was (**Figure 1b**) significant different among three groups. For PFHxS and PFOS (**Figure 1c** and **1d**), there were no significant differences among the three groups. But the PFAS concentration for the firefighters is more varied which may be attributed to the more frequent exposure to PFHxS and PFOS in AFFF, as these chemicals are components of AFFF.

**Figure 1.**
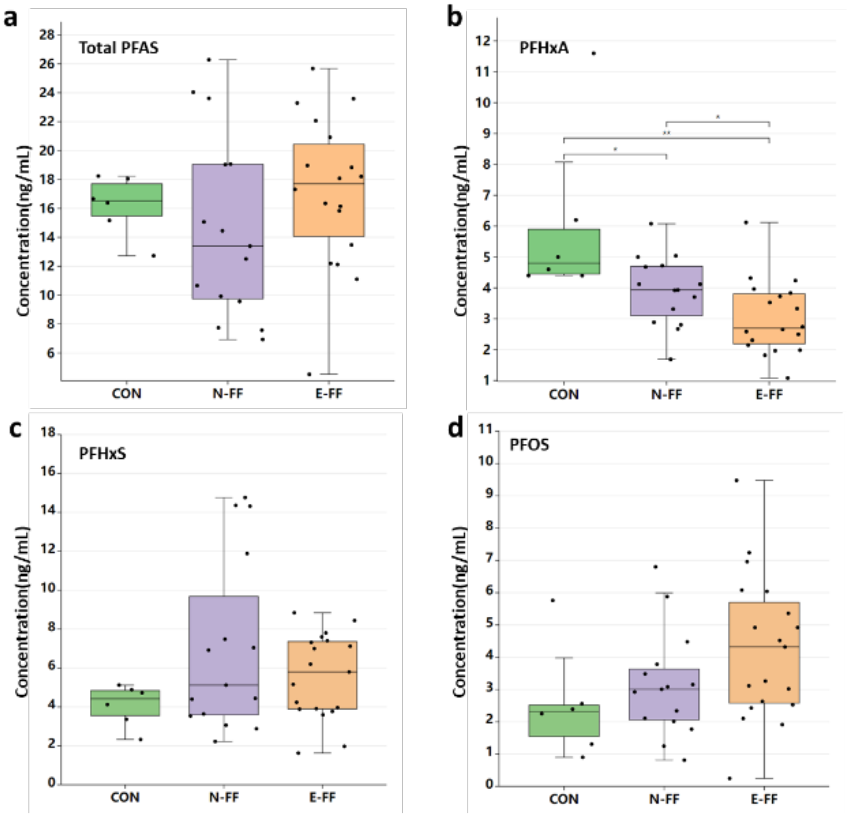
Total PFAS (a), PFHxA (b), PFHxS (c), PFOS (d) plasma concentration of control group (CON), N-FF and E-FF.

### 3.3 MiRNA Sequencing of Plasma Exosomes

According to the results of PFAS level, a total of 10 samples (three CONs, three N-FFs and four E-FFs) are selected and sent for miRNA sequencing. Exosomes from plasma were extracted and characterized (Figure S1). The flat round structure of exosomes can be clearly observed under the transmission electron microscope (TEM). Nanoparticle tracking analysis (NTA) and dynamic light scattering (DLS) shows that the diameter is mostly distributed around 70 nm, and the particle size is in line with the size distribution range of exosomes.

After assessing the sequencing read quality using miRTrace (Supplementary Table S2), we decided to exclude the outlier sample 38E due to significantly lower miRNA reads. Both the multi-dimensional scaling (MDS) plot and the heatmap (**Figure 2**) visually demonstrate substantial dispersion and dissimilarity within each group. Consequently, we opted not to proceed with further differential expression analysis.

**Figure 2.**
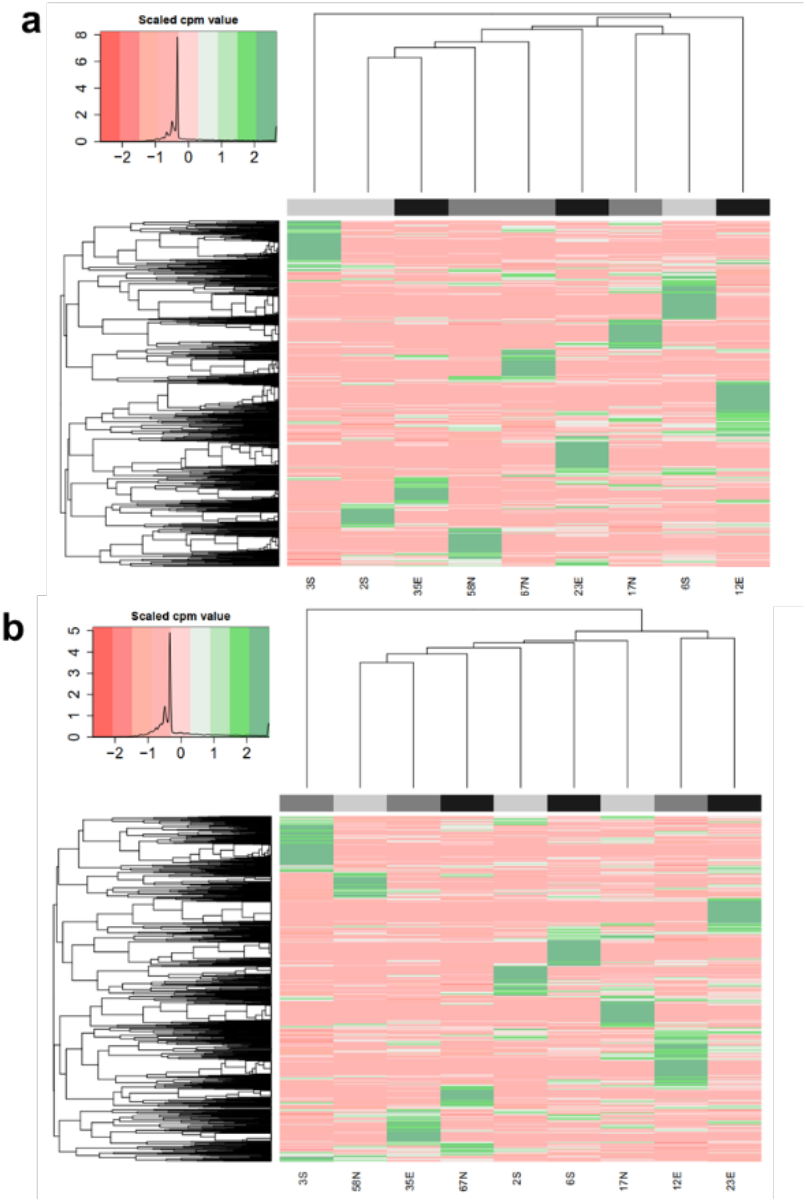
Heatmap of firefighters’ miRNA-hairpin (a), Heatmap of firefighters’ miRNA-mature (b).

### 3.4 Population characteristics of NHANES population

During the 2017-2018 period, data on serum PFAS levels was available for only one-third of the survey population, and a summary of their main characteristics is presented in **Table 2**. The average age of NHANES participants with PFAS data was 45.26, which is higher than the average age of the overall survey population (34.4). This age difference is primarily due to the eligibility criteria for PFAS testing, which required participants to be 12 years of age or older. Across other covariates, we observed similar distributions within their respective categories. In our sample, NHANES participants were predominantly Non-Hispanic White (34.6%), possessed education levels higher than college (56.2%), reported a family income-to-poverty ratio between 1 and 3 (43.9%), had a BMI exceeding 30 (38.7%), identified as non-smokers (59.7%), were moderate drinkers (69.5%), and engaged in low levels of physical activity (37.9%). In terms of self-reported health conditions, more than 30% of the population in our sample had hypertension or cardiovascular-related diseases.

### 3.5 Comparison with PFAS Levels in NHANES

Additionally, we conducted a Mann-Whitney U test, given the skewed distribution of PFAS levels, to compare PFAS levels between firefighters and the general population data from NHANES. Our analysis did not reveal any statistically significant differences in PFAS levels between these two populations.

### 3.6 PFAS Level and Biochemical Measure in NHANES

Initially, we examined the impact of increasing log-transformed total serum PFAS levels on various biochemical indicators through survey-weighted multivariate linear regression analysis. Our findings indicated that for each 1-unit increase in log-transformed weighted total PFAS concentration, there were positive associations with specific biochemical indicators, including a 0.17 µg/L increase in ferritin, a 0.02 µg/mL increase in serum iron, a 0.012 mg/dL increase in fasting glucose, a 0.04 mg/dL increase in cholesterol, a 0.05 U/L increase in serum ALT, a 0.01 g/dL increase in serum albumin, a 0.02 IU/L increase in serum ALP, a 0.04 U/L increase in serum AST, a 0.005 mmol/L increase in serum bicarbonate, a 0.0007 mmol/L increase in serum chloride, a 0.0002 mmol/L increase in serum sodium, a 0.04 mg/dL increase in serum total bilirubin, a 0.005 mg/dL increase in serum total calcium, a 0.0035 g/dL increase in total protein, and a 0.0031 mg/dL increase in serum uric acid. Conversely, urine albumin, urine creatinine, serum UIBC, serum glycohemoglobin, RBC folate, triglycerides, serum blood urea nitrogen, serum creatinine, serum phosphorus, serum potassium exhibited negative associations with log transformed weighted total PFAS levels (**Table 3**).

**Table 3.** Difference in continuous biochemical indicators and the relative odds ratio (95%) according to serum PFAS levels among participants in NAHANES 2017-2018.

|  | Linear model |  | Multinomial model 1 | Multinomial model 2 |
| --- | --- | --- | --- | --- |
| Biochemical indicators | Coefficient (95% CI) | Categories | Coefficient (95% CI) |  |
| Albumin_urine (μg/mL) | -0.1259 (-0.2154, -0.0363) | Microalbuminuria | 1.00000 (1.00000, 1.00000) | 1.00000 (1.00000, 1.00000) |
|  |  | Macroalbuminuria | 1.00000 (1.00000, 1.00000) | 1.00000 (1.00000, 1.00000) |
| Creatinine_urine (mg/dL) | -0.0380 (-0.0901, 0.0142) | Low | 1.00000 (1.00000, 1.00000) | 1.00000 (1.00000, 1.00000) |
|  |  | High | 1.00000 (1.00000, 1.00000) | 1.00000 (1.00000, 1.00000) |
| Ferritin (ug/L) | 0.1679 ((0.1082 0.2275)) | Low | 1.00000 (1.00000, 1.00000) | 1.00000 (1.00000, 1.00000) |
|  |  | High | 1.00000 (1.00000, 1.00000) | 1.00000 (1.00000, 1.00000) |
| Iron_serum (µg/mL) | 0.0196 (-0.0088, 0.0479) | Low | 1.00000 (1.00000, 1.00000) | 1.00000 (1.00000, 1.00000) |
|  |  | High | 1.00000 (1.00000, 1.00000) | 1.00000 (1.00000, 1.00000) |
| UIBC_serum (µg/dL) | -0.0061 (-0.024,0.0118) | Low | 1.00000 (1.00000, 1.00000) | 1.00000 (1.00000, 1.00000) |
|  |  | High | 1.00000 (1.00000, 1.00000) | 1.00000 (0.99999, 1.00000) |
| Glycohemoglobin_serum | -0.0001 (-0.0076, 0.0075) | Prediabetes | 1.00000 (1.00000, 1.00000) | 1.00000 (1.00000, 1.00000) |
|  |  | Diabetes | 1.00000 (1.00000, 1.00000) | 1.00000 (1.00000, 1.00000) |
| Folate_RBC (ng/mL) | -0.0236 (-0.0748, 0.0277) | Low | 1.00000 (1.00000, 1.00000) | 1.00000 (1.00000, 1.00000) |
|  |  | High | 1.00000 (1.00000, 1.00000) | 1.00000 (1.00000, 1.00000) |
| Fasting_glucose (mg/dL) | 0.0121(-0.0057, 0.0299) | Prediabetes | 1.00000 (1.00000, 1.00000) | 1.00000 (1.00000, 1.00000) |
|  |  | Diabetes | 1.00000 (1.00000, 1.00000) | 1.00000 (1.00000, 1.00000) |
| Cholesterol (mg/dL) | 0.0376 [(0.0224, 0.0527] | Low | 1.00000 (1.00000, 1.00000) | 1.00000 (1.00000, 1.00000) |
| Triglyceride (mg/dL) | -0.0111(-0.0692, 0.0469) | Low | 1.00000 (1.00000, 1.00000) | 1.00000 (1.00000, 1.00000) |
| Serum_ALT (U/L) | 0.0464 (0.0123, 0.0805) | Mildly high | 1.00000 (1.00000, 1.00000) | 1.00000 (1.00000, 1.00000) |
|  |  | High | 1.00000 (1.00000, 1.00000) | 1.00000 (1.00000, 1.00000) |
| Albumin_serum (g/dL) | 0.0118 (0.0065, 0.0171) | Low | 1.00000 (1.00000, 1.00000) | 1.00000 (1.00000, 1.00000) |
| ALP_serum (IU/L) | 0.0237 (0.0011, 0.0463) | Low | 1.00000 (1.00000, 1.00000) | 1.00000 (1.00000, 1.00000) |
|  |  | High | 1.00000 (1.00000, 1.00000) | 1.00000 (1.00000, 1.00000) |
| AST_serum (U/L) | 0.0356 (0.0115, 0.0597) | Low | 1.00000 (1.00000, 1.00000) | 1.00000 (1.00000, 1.00000) |
|  |  | High | 1.00000 (1.00000, 1.00000) | 1.00000 (1.00000, 1.00000) |
| Bicarbonate_serum (mmol/L) | 0.0052 (-0.0019, 0.0123) | Low | 1.00000 (1.00000, 1.00000) | 1.00000 (1.00000, 1.00000) |
|  |  | High | 1.00000 (1.00000, 1.00000) | 1.00000 (1.00000, 1.00000) |
| Blood_urea_nitrogen_serum (mg/dL) | -0.0152 (-0.0374, 0.0070) | Low | 0.99999 (0.99999, 0.99999) | 1.00000 (1.00000, 1.00000) |
|  |  | High | 1.00000 (1.00000, 1.00000) | 1.00000 (1.00000, 1.00000) |
| Chloride_serum (mmol/L) | 0.0007 (-0.0013, 0.0028) | Low | 1.00000 (1.00000, 1.00000) | 1.00000 (1.00000, 1.00000) |
|  |  | High | 1.00000 (1.00000, 1.00000) | 1.00000 (1.00000, 1.00000) |
| Creatinine_serum (mg/dL) | -0.0367(-0.0529, -0.0206) | Low | 1.00000 (1.00000, 1.00000) | 1.00000 (1.00000, 1.00000) |
|  |  | High | 1.00000 (1.00000, 1.00000) | 1.00000 (1.00000, 1.00000) |
| Phosphorus_serum (mg/dL) | -0.0019 (-0.0128, 0.008) | Low | 1.00000 (1.00000, 1.00000) | 1.00000 (1.00000, 1.00000) |
|  |  | High | 1.00000 (1.00000, 1.00000) | 1.00000 (1.00000, 1.00000) |
| Potassium_serum(mmol/L) | -0.0042 (-0.0106, 0.0023) | Low | 1.00000 (1.00000, 1.00000) | 1.00000 (1.00000, 1.00000) |
|  |  | High | 1.00000 (1.00000, 1.00000) | 1.00000 (1.00000, 1.00000) |
| Sodium_serum (mmol/L) | 0.0002 (-0.0012, 0.0017) | Low | 1.00000 (1.00000, 1.00000) | 1.00000 (1.00000, 1.00000) |
|  |  | High | 1.00000 (1.00000, 1.00000) | 1.00000 (1.00000, 1.00000) |
| Total_bilirubin_serum (mg/dL) | 0.0378 (0.0037, 0.0720) | Low | 1.00000 (1.00000, 1.00000) | 1.00000 (1.00000, 1.00000) |
| Total_calcium_serum(mg/dL) | 0.0046 (0.0018, 0.0074) | Low | 1.00000 (1.00000, 1.00000) | 1.00000 (1.00000, 1.00000) |
|  |  | High | 1.00000 (1.00000, 1.00000) | 1.00000 (1.00000, 1.00000) |
| Total_protein_serum(g/dL) | 0.0035 (-0.0006, 0.0075) | Low | 1.00000 (1.00000, 1.00000) | 1.00000 (1.00000, 1.00000) |
|  |  | High | 1.00000 (1.00000, 1.00000) | 1.00000 (1.00000, 1.00000) |
| Uric_acid_serum (mg/dL) | 0.0031 (-0.0140, 0.0202) | Low | 1.00000 (1.00000, 1.00000) | 1.00000 (1.00000, 1.00000) |
|  |  | High | 1.00000 (1.00000, 1.00000) | 1.00000 (1.00000, 1.00000) |

Subsequently, we conducted multinomial regression analysis to investigate the relationship between serum PFAS levels and categories of biochemical indicators by multinomial models. The majority of the exponentiated coefficients were close to 1.0, indicating nearly equal odds when compared to the reference (normal range). This suggests that serum PFAS levels were not significant predictors for any of the biochemical indicator outcomes (**Table 3**; Supplementary Table S5; Supplementary Table S6).

## 4. Discussion

The objective of the study is to investigate the potential mechanistic links between PFAS exposure and cancer biomarkers among firefighters. In recent years, exosomes have emerged as a fascinating area of study. The potential role of exosomal miRNAs in mediating the effects of PFAS exposure on cancer risk is an intriguing hypothesis. While existing studies have predominantly explored the association between PFAS levels in firefighters’ blood and cancer risk, only a few have ventured into mechanistic evaluations^29^. In this study, we take this exploration a step further by evaluating the miRNA content of exosomes in conjunction with PFAS levels in the blood of firefighters. Our objectives include not only quantifying PFAS and metabolites in the blood samples of differentially exposed firefighter populations but also conducting RNA-Seq and informatics to establish the relationship between PFAS levels in plasma and miRNA levels in exosomes. These findings were compared with the general population data from NHANES to provide context for the results.

The study provided a comprehensive overview of the characteristics of firefighters, highlighting key differences between new firefighters (N-FF) and experienced firefighters (E-FF). The analysis revealed that there were no statistically significant differences in total PFAS levels in plasma between the general population, new firefighters (N-FF), and experienced fire-fighters (E-FF). Most participants in all groups had total PFAS values falling within the 10-30 ng/mL range. The average experience of E-FFs was significantly higher than that of N-FFs, which could influence their exposure to PFAS. New firefighters, with less experience, may not have encountered as many PFAS sources or situations, thus contributing to lower levels of certain PFAS compounds. Experienced firefighters have encountered a wider range of incidents involving PFAS-containing materials and AFFF, potentially explaining the observed higher PFAS levels, particularly for compounds like PFOS. No significant differences were found for PFHxS levels among the three groups, suggesting consistent exposure patterns. The findings regarding PFAS levels in firefighters align with some previous studies, which have also reported elevated PFAS levels among firefighting personnel due to the use of AFFF containing these chemicals^7,9^. The lack of significant differences in PFHxS and PFOS levels may reflect the prevalence of these compounds in AFFF. While our results show consistency with certain existing research^43-45^, PFHxA is most abundant in the control group, which may be due to the fact that it is a product of breakdown of other PFAS and is widely present in the environment. As a result, it can be exposed to a variety of routes, including air inhalation, dietary intake, and skin exposure to PFHxA-containing products. The significant difference in PFHxA levels in the samples collected requires further exploration.

The miRNA sequencing of plasma exosomes in the study aimed to uncover potential mechanistic insights into PFAS exposure and its effects on firefighters. The presence of substantial dispersion and dissimilarity within each group, as indicated by the MDS plot and heatmap, suggests considerable heterogeneity in miRNA expression. This may be attributed to individual variability in miRNA expression or other confounding factors not addressed in our study^29^. While the analysis did not proceed to differential expression analysis, the variations in miRNA profiles among different groups, even within the same occupation, emphasize the complexity of the biological responses to PFAS exposure.

The results were compared with the NHANES data for PFAS levels^8,46-48^. The analysis did not reveal any statistically significant differences in PFAS levels between firefighters and the general population. This suggests that total PFAS exposure patterns in firefighters may not substantially differ from the general population, at least in terms of PFAS levels in plasma^49,50^. This outcome also points to the ubiquitous exposure to PFAS for all citizens. It is likely that the general population is exposed to PFAS through a variety of sources, including consumer products, food, and environmental contamination^1^. This widespread exposure may reduce the relative difference in PFAS levels between firefighters and the general population. Additionally, the eligibility criteria for PFAS testing in NHANES, was 12 years of age or older^51^, may have introduced age-related differences between the two groups. While not statistically significant, the trends in PFHxS and PFOS for new and experienced firefighters is worth further exploration for mechanistical causes of increased rates of cancer incidence in the firefighting community.

In the investigation of the relationship between log-transformed total serum PFAS levels and various biochemical indicators, some positive associations with specific indicators were found, such as ferritin, serum iron, fasting glucose, cholesterol, and others. Conversely, some indicators exhibited negative associations. Multinomial regression analysis, however, indicated that serum PFAS levels were not significant predictors for any of the biochemical indicator outcomes.

Several limitations must be acknowledged in the study. One of the primary limitations of this study is the relatively small sample size, which included 34 firefighter participants. This limited sample size may not fully represent the diverse population of firefighters, and the findings may not be generalizable to the entire firefighting community. The study’s recruitment of participants from a single community may introduce selection bias. Firefighters from different regions or with varying exposure scenarios may exhibit different PFAS levels and health outcomes. The study primarily assessed PFAS exposure through plasma levels, but did not explore potential dermal, inhalation, or dietary exposure pathways, which may contribute to overall exposure patterns among firefighters. Therefore, the findings may not fully reflect the broader firefighter population. The study utilized a cross-sectional design to assess PFAS exposure and health indicators at a single point in time. Cross-sectional studies do not establish causality or account for changes over time. Longitudinal studies may offer more insights into the temporal relationship between PFAS exposure and health outcomes. While the study included miRNA sequencing of plasma exosomes, we did not extend further to differential expression analysis due to data quality concerns and the exclusion of outliers. This limitation restricts the assessment of potential mechanistic insights into how PFAS exposure affects firefighters’ health at the molecular level.

Despite these limitations, the study provides valuable insights into the PFAS exposure of firefighters and its potential impact on biochemical indicators. While the results did not demonstrate substantial differences in PFAS levels between firefighters and the general population, the unique exposure patterns, and the potential impact of specific PFAS compounds, highlight the complexity of this issue. Further research is needed to explore the long-term health effects of PFAS exposure in firefighting personnel and identify effective mitigation strategies.

## 5. Conclusion

This study provides valuable insights into the characteristics of firefighters, their PFAS levels, and the potential impact of PFAS on their health. Firefighters, both new and experienced, displayed consistent sociodemographic and lifestyle characteristics, with notable differences in work experience. While the total PFAS levels in plasma did not significantly differ between firefighters and the general population, variations in individual PFAS compounds were observed. The study’s findings suggest substantial heterogeneity in miRNA expression among the study participants. The comparison with the general population did not reveal significant differences in PFAS levels, possibly due to widespread environmental PFAS exposure.

## AUTHOR INFORMATION

### Author Contributions

Conceptualization, J.I. and X.Z.; methodology, all authors.; software, M.S, C.F; validation, X.Z, MS; data curation, M.S, X.Z.; writing—original draft preparation, X.Z. and M.S; writing—review and editing, all authors; supervision, J.I.; project administration, J.I, and MJS. All authors have read and agreed to the published version of the manuscript. ^#^These authors contributed equally.

## ACKNOWLEDGMENT

We thank Alexander Ulanov of the Roy J. Carver Biotechnology Center Metabolomics Core for PFAS analysis. Steve Doggett, from IFSI, is acknowledged for facilitating the recruitment of firefighters in this study. Tumor Engineering and Phenotyping facility of the Cancer Center at Illinois is acknowledged for help in exosome experiments.

