## Supplemental Information for "MicroRNAs and PFAS: A Pilot Study in Blood Collected from Firefighters"

### SUPPLEMENTARY MATERIALS

### Contents list

S.1 Chemicals and reagents
S.2 Quantification of PFAS in Blood
S.3 Quality assurance and quality control (QA/QC)
Figure S1 and Table S1-S3
S.4 Health Physical Activity Inventory

*S.1 Chemicals and reagents*

MPFAC-24ES, a mixture containing 19 mass-labelled standards containing $C_4$-$C_{14}$ PFCAs, $C_4$-$C_{10}$ PFSAs, FOSA, N-MeFOSAA, N-EtFOSAA, as well as 4:2, 6:2, and 8:2FTS, was purchased from Wellington Laboratories (Guelph, ON, Canada). Target compounds included 20 PFAS contained within the PFAC-24PAR mixture purchased from Wellington Laboratories (Canada), as well as an additional PFAS purchased from SynQuest Laboratories (USA). The stock solutions were prepared in methanol and kept in polypropylene (PP) tubes at 4°C. HPLC-grade methanol, with a purity of at least 99.9%, from Sigma-Aldrich (USA), was used as a solvent for HPLC analysis.

*S.2 Quantification of PFAS in Blood*

Plasma samples were analyzed for PFAS metabolites using liquid chromatography-mass spectrometry (LC-MS) by the Carver Metabolomics Core of the Roy J. Carver Biotechnology Center, University of Illinois Urbana-Champaign. Briefly, 50 µL plasma were added to 2 mL polypropylene vials, followed by 5 µL of a mixture of 19 stable isotope- and deuterium-labeled internal standards and 200 µL methanol. Upon vortexing and centrifugation, 150 µL supernatant was transferred to polypropylene HPLC vials. Using an Agilent 1260 Infinity II HPLC system (Agilent Technologies, Santa Clara, CA, USA), 10 µL were injected and metabolites separated using a Phenomenex Kinetex PS C18 100A (2.6µm, 100 x 4.6mm) column (Phenomenex, Torrance, CA, USA) via a gradient method consisting of mobile phase A: H2O+20mM ammonium acetate and mobile phase B: methanol running at a flow rate of 0.35 mL/min. The gradient was 0-2 min = 90% A; 2-10min = 0% A; 18.1-24min = 90% A. A Sciex 6500+ triple quadrupole MS (Sciex, Framingham, MA, USA) operating in negative ionization mode used multiple reaction monitoring (MRM) to screen for 21 PFAS metabolites. Metabolites were quantified with Sciex Analyst software using 5- to 7-point calibration curves adjusted for corresponding stable isotope- and deuterium-labeled internal standards. A delay column (Halo PFAS delay column, 2.7 µm, 4.6 mm x 50 mm; Advanced Materials Technology, Wilmington, DE, USA) was placed between the mobile phase mixer and sample injector to momentarily trap any system related PFAS interferences due to tubing or mobile phase.

*S.3 Quality assurance and quality control (QA/QC)*

. Metabolites were quantified with Sciex Analyst software using 5- to 7-point calibration curves adjusted for corresponding stable isotope- and deuterium-labeled internal standards. A delay column (Halo PFAS delay column, 2.7 µm, 4.6 mm x 50 mm; Advanced Materials Technology, Wilmington, DE, USA) was placed between the mobile phase mixer and sample injector to momentarily trap any system related PFAS interferences due to tubing or mobile phase. Choice of materials used throughout sample processing and analysis sought was done with the intent of minimizing PFAS contamination. Blanks were periodically analyzed throughout the sequence to test for the presence of PFAS contamination. QCs were run to assess reproducibility of quantitation at varying concentrations. The analytical lab was blinded to the identity of the sample groups. All samples were randomized and analyzed in a single batch.

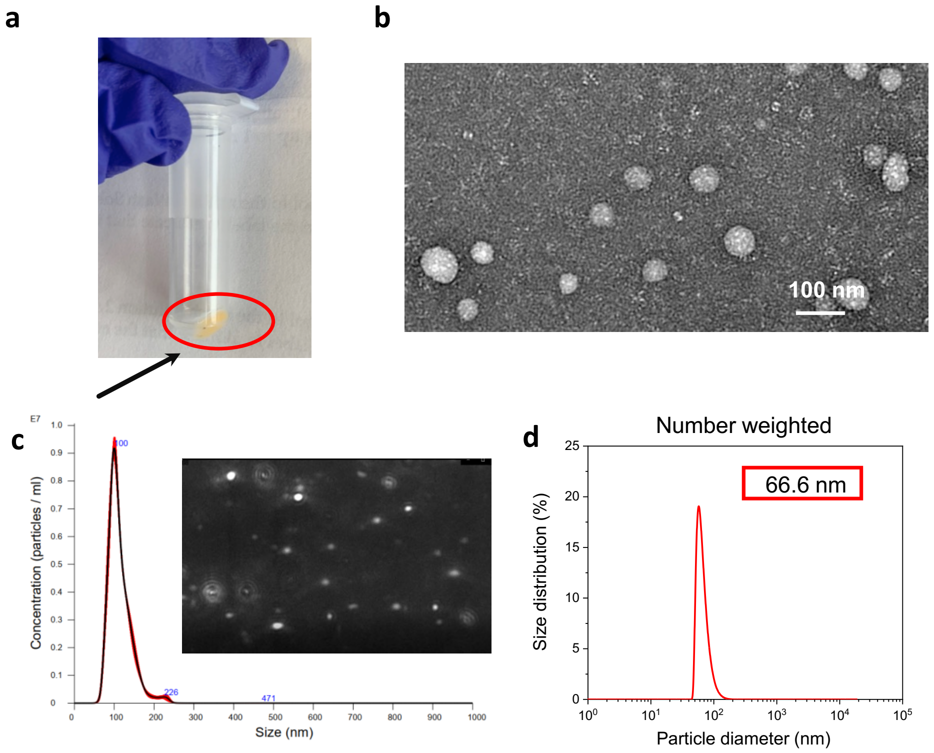


**Figure S1, (a)**. Exosomes in plasma; **(b)**. Morphology of plasma exosomes under TEM; **(c)**. NTA of plasma exosomes; **(d)**. Particle size of plasma exosomes.

**Supplementary table 1. Characteristics of firefighters**

| Sample | Age | Gender | Weight (kg) | Height (m) | BMI | Drinks Alcohol | Taking Medications | Days of Strenuous Exercise | Years in Service | Working Fire Responses (Past Year) |
|---|---|---|---|---|---|---|---|---|---|---|
| 17N | 23 | M | 77.11 | 1.91 | 21.25 | Y | Y | 4 | 2 | 4-5 |
| 40N | 20 | M | 88.45 | 1.93 | 23.74 | Y | N | 5 | 3 | 115 |
| 45N | 25 | M | 86.18 | 1.91 | 23.75 | Y | N | 3 | 0 | 0 |
| 47N | 22 | M | 97.52 | 1.83 | 29.16 | N | N | 4 | 3 | 10 |
| 52N | 25 | M | 86.18 | 1.75 | 28.06 | Y | N | 3 | 3 | 4 |
| 54N | 21 | M | 79.38 | 1.74 | 26.22 | Y | N | 3 | 0 | 0 |
| 55N | 24 | M | 81.65 | 1.78 | 25.83 | Y | N | 2 | 0 | 0 |
| 57N | 27 | M | 108.86 | 1.65 | 39.94 | Y | Y | 2 | 0.5 | 2 |
| 58N | 31 | M | 82.10 | 1.75 | 26.73 | Y | N | 5 | 2.5 | 3 |
| 59N | 30 | M | 90.72 | 1.83 | 27.12 | Y | N | 5 | 1.5 | 2 |
| 61N | 32 | F | 50.35 | 1.55 | 20.97 | Y | N | 5 | 0 | 3 |
| 64N | 24 | M | 86.18 | 1.80 | 26.50 | N | Y | 4 | 3 | 15 |
| 65N | 28 | F | 64.86 | 1.73 | 21.74 | N | Y | 5 | 0 | 1 |
| 67N | 29 | M | 77.11 | 1.78 | 24.39 | N | Y | 5 | 0 | 0 |
| 68N | 26 | M | 113.40 | 2.01 | 28.16 | N | Y | 5 | 2 | 2 |
| 12E | 36 | M | 111.13 | 1.75 | 36.18 | N | Y | 3 | 17 | 8 |
| 18E | 51 | M | 127.01 | 1.83 | 37.97 | Y | Y | 1 | 17 | 1 |
| 20E | 32 | M | 124.74 | 1.75 | 40.61 | N | Y | 5 | 17 | 15 |
| 22E | 44 | M | 75.75 | 1.83 | 22.65 | Y | N | 5 | 25 | 15 |
| 23E | 41 | M | 90.72 | 1.83 | 27.12 | Y | N | 3 | 24 | 45 |
| 24E | 43 | M | 108.86 | 1.83 | 32.55 | Y | Y | 4 | 25 | 10 |
| 29E | 43 | M | 149.69 | 1.93 | 40.17 | Y | Y | 3 | 26 | 5 |
| 31E | 39 | M | 127.01 | 1.91 | 35.00 | Y | Y | 0 | 16 | 12 |
| 32E | 37 | M | 95.25 | 1.80 | 29.29 | Y | N | 4 | 15 | 6 |
| 33E | 53 | M | 138.35 | 1.83 | 41.37 | Y | Y | 2 | 30 | 0 |
| 34E | 43 | M | 74.84 | 1.73 | 25.09 | N | N | 2 | 26 | 4 |
| 35E | 47 | M | 63.50 | 1.83 | 18.99 | Y | N | 1 | 21 | 7 |
| 36E | 45 | M | 133.81 | 1.88 | 37.88 | Y | Y | 2 | 30 | 6 |
| 37E | 51 | M | 92.99 | 1.88 | 26.32 | N | N | 1 | 32 | 5 |
| 38E | 37 | M | 92.99 | 1.78 | 29.41 | Y | Y | 1 | 15 | 6 |
| 39E | 53 | M | 99.79 | 1.73 | 33.45 | Y | N | 5 | 30 | 15 |

| | | | | | | | | | | |
|---|---|---|---|---|---|---|---|---|---|---|
| 43E | 46 | M | 74.84 | 1.75 | 24.37 | Y | N | 2 | 20 | 20 |
| 60E | 52 | M | 106.59 | 1.83 | 31.87 | Y | Y | 3 | 21 | 10 |
| 62E | 36 | M | 72.57 | 1.75 | 23.63 | Y | N | 7 | 20 | 10 |

**Supplementary table 2. Descriptive statistics before and after miRTrace quality control**

| Sample | Total reads | QC-passed reads | miRNA reads |
|---|---|---|---|
| 2S | 27,746,244 | 27,586,674 | 9,752,977 |
| 35E | 30,656,625 | 30,494,280 | 7,331,531 |
| 23E | 30,816,067 | 30,721,960 | 4,682,025 |
| 67N | 25,685,839 | 25,362,176 | 10,066,308 |
| 17N | 33,191,655 | 33,009,001 | 10,015,862 |
| 3S | 32,000,528 | 31,541,637 | 5,201,497 |
| 6S | 30,424,651 | 30,228,999 | 14,242,280 |
| 58N | 27,975,105 | 27,921,488 | 1,845,735 |
| 38E | 38,142,391 | 37,364,129 | 14,824 |
| 12E | 33,871,686 | 33,579,028 | 6,223,323 |

**Supplementary table 3. Full Biochemical Catogories**

| | | |
|---|---|---|
| Biochemical Group | | |
| Albumin_urine (μg/mL) | | |
| Creatinine_urine (mg/dL) | Female | Male |
| Ferritin (ug/L) | Female | Male |
| Iron_serum (μg/mL) | Female | Male |
| UIBC_serum (μg/dL) | | |
| Glycohemoglobin_serum | | |
| folate_RBC (ng/mL) | | |
| Fasting_glucose (mg/dL) | | |
| Cholesterol (mg/dL) | | |
| Triglyceride (mg/dL) | | |
| Vitamin_D (nmol/L) | | |
| Serum_ALT (U/L) | | |
| Albumin_serum (g/dL) | | |
| ALP_serum (IU/L) | | |
| AST_serum (U/L) | | |
| Bicarbonate_serum (mmol/L) | | |
| Blood_urea_nitrogen_serum (mg/dL) | | |
| Chloride_serum (mmol/L) | | |
| Creatinine_serum (mg/dL) | Female | Male |

*S.4 Health Physical Activity Inventory*

Please complete the survey below.

Thank you!

### Physiological Characteristics

Age

______________________________

Gender

◯ Male
◯ Female

What is your current weight?

______________________________
(lbs)

What is your current height?

______________________________
(inches)

What is your current waist circumference?

______________________________
(in)

Have you traveled outside the United States in the past month?

◯ Yes
◯ No

Did you have COVID in the past month?

◯ Yes
◯ No

Are you a smoker or tobacco user?

◯ Yes
◯ No

On average, how many packs of cigarettes do you smoke per day?

◯ 0
◯ 1
◯ 2
◯ 3
◯ 4+

On average, how many packs of cigarettes do you smoke per week?

◯ 0
◯ 1
◯ 2
◯ 3
◯ 4+

Do you drink alcohol?

◯ Yes
◯ No

On average, how many servings of alcohol per week do you consume?

◯ 0
◯ 1
◯ 2
◯ 3
◯ 4+

Are you currently taking any medications or antibiotics?

◯ Yes
◯ No

| | |
|---|---|
| If yes, please describe. | ______________________ |
| Do you have a family history of cardiovascular disease (CVD)? | ◯ Yes<br>◯ No |
| Do you have any known CVD? | ◯ Yes<br>◯ No |
| Please describe your CVD. | ______________________ |
| Do you have any known neurological diseases? | ◯ Yes<br>◯ No |
| Please describe your neurological disease. | ______________________ |
| Do you have any known digestive disorders including abnormalities with swallowing, esophageal or bowel strictures, gastrointestinal obstruction or fistulas? | ◯ Yes<br>◯ No |
| Please describe your digestive disorder. | ______________________ |

### Physical Activities

| | |
|---|---|
| How would you describe your overall fitness?<br>*Note average is work out 3 times a week for 30 min. or more per session. | ◯ Poor<br>◯ Below average<br>◯ Average<br>◯ Very good<br>◯ Excellent |
| In the last 3 months, how many days a week have you spent 30 minutes or more in moderate to strenuous exercise? | ◯ 0<br>◯ 1<br>◯ 2<br>◯ 3<br>◯ 4<br>◯ 5<br>◯ 6<br>◯ 7<br>(days) |

### Typical PPE Usage, Cleaning, and Exposures

| | |
|---|---|
| During overhaul, do you ever wear your SCBA? | ◯ Yes<br>◯ No |
| What percent of time during overhaul do you usually wear your SCBA? | ______________________<br>(%) |

**What PPE do you typically wear 100% of the time when involved in the following activities and types of fires? (Check all that apply)**

| | Structural knockdown | Structural overhaul | Roof ventilation | Vehicle fire knockdown | Vehicle fire overhaul | Vegetation fires |
|---|---|---|---|---|---|---|
| Fire fighting turnout coat/pants | ☐ | ☐ | ☐ | ☐ | ☐ | ☐ |
| Fire fighting boots | ☐ | ☐ | ☐ | ☐ | ☐ | ☐ |
| SCBA and on air | ☐ | ☐ | ☐ | ☐ | ☐ | ☐ |
| Fire fighting helmet | ☐ | ☐ | ☐ | ☐ | ☐ | ☐ |
| Fire fighting hood | ☐ | ☐ | ☐ | ☐ | ☐ | ☐ |
| Fire fighting gloves | ☐ | ☐ | ☐ | ☐ | ☐ | ☐ |
| Work gloves | ☐ | ☐ | ☐ | ☐ | ☐ | ☐ |
| Eye/face protection other than SCBA face piece | ☐ | ☐ | ☐ | ☐ | ☐ | ☐ |

**What PPE do you typically wear 50% or more when involved in the following activities and types of fires? (Check all that apply)**

| | Structural knockdown | Structural overhaul | Roof ventilation | Vehicle fire knockdown | Vehicle fire overhaul | Vegetation fires |
|---|---|---|---|---|---|---|
| Fire fighting turnout coat/pants | ☐ | ☐ | ☐ | ☐ | ☐ | ☐ |
| Fire fighting boots | ☐ | ☐ | ☐ | ☐ | ☐ | ☐ |
| SCBA and on air | ☐ | ☐ | ☐ | ☐ | ☐ | ☐ |
| Fire fighting helmet | ☐ | ☐ | ☐ | ☐ | ☐ | ☐ |
| Fire fighting hood | ☐ | ☐ | ☐ | ☐ | ☐ | ☐ |
| Fire fighting gloves | ☐ | ☐ | ☐ | ☐ | ☐ | ☐ |
| Work gloves | ☐ | ☐ | ☐ | ☐ | ☐ | ☐ |
| Eye/face protection other than SCBA face piece | ☐ | ☐ | ☐ | ☐ | ☐ | ☐ |

On average, how often each year is your turnout gear laundered?

- ○ After each activity
- ○ Between 2-5 activities
- ○ Weekly
- ○ Monthly
- ○ Every 2-3 months
- ○ Yearly

(times)

On average, how often do you perform gross decon on your turnout gear?

- ○ After each activity
- ○ Between 2-5 activities
- ○ Weekly
- ○ Monthly
- ○ Every 2-3 months
- ○ Yearly

(times)

| | |
|---|---|
| On average, how often is your hood laundered? | ◯ After each activity<br>◯ Between 2-5 activities<br>◯ Weekly<br>◯ Monthly<br>◯ Every 2-3 months<br>◯ Yearly<br>(times) |
| Where do you store gear that has been used in a response? (Check all that apply) | ☐ At the fire station in the gear storage area<br>☐ At the fire station in living areas<br>☐ In the fire apparatus<br>☐ In my personal vehicle<br>☐ In my home<br>☐ Other |
| Please describe where else you store gear that has been used in a response. | ______________________ |
| After fire suppression activities, do you notice soot on your skin after removing your SCBA or turnout gear? | ◯ Yes<br>◯ No |
| If yes, which body part(s) do you most often find soot? (Check all that apply). | ☐ Hands<br>☐ Arms<br>☐ Neck<br>☐ Face<br>☐ Chest<br>☐ Abdomen<br>☐ Groin<br>☐ Legs/Feet |
| Do you usually shower after a shift that included fire suppression actitivites? | ◯ Yes<br>◯ No |

### Fire Service History

| | |
|---|---|
| How many total years have you served in the fire service? | ______________________<br>(years) |
| What is your current rank/title? | ______________________ |
| In the past year, approximately how many working fire responses (structure, vehicle, wildland, etc.) were you exposed to smoke and other products of combustion? | ______________________ |
| Throughout your fire service career which type of fuels and chemicals were you exposed to? (Check all that apply.) | ☐ Pallet & straw/hay/excelsior<br>☐ OSB or other manufactured wood products<br>☐ Theatrical smoke<br>☐ Gas fired burner<br>☐ Theatrical smoke & gas burners combined<br>☐ Firefighting foam<br>☐ Other (describe below) |
| Other (please describe) | ______________________ |
